# Estimation of Location-specific Variation in Regional Food Web Interactions

**DOI:** 10.1101/2024.01.17.575986

**Authors:** Aaron Brothers, Vinicio Santillán, Marta Quitián, Eike Lena Neuschulz, Matthias Schleuning, Katrin Böhning-Gaese, Orlando Alvarez, Christopher Thron

## Abstract

Food-web interactions are commonly aggregated across sampling locations and represented by a single interaction strength for an entire region. This practice implicitly treats interaction strengths as fixed regional constants, even though ecological interactions may vary substantially among locations because of local environmental conditions, species abundances, and habitat structure. We argue that food-web interaction strengths should instead be viewed as spatially distributed latent variables whose regional distributions must be inferred from observations collected at a limited number of sites.

To estimate this spatial variation, we develop a Bayesian framework based on transect observations from multiple locations within a region. Species–species interaction measurements at each location are modeled as Poisson random variables whose means are proportional to local interaction strengths. Preliminary analyses suggest that location-dependent interaction strengths may be simply approximated by a uniform distribution on an interval. Under this assumption, we estimate the endpoints of the interval and thereby characterize both the mean interaction strength and its spatial variability. For the case of three sampling locations, we derive graphical representations of the resulting estimates and their uncertainties. Among other results, we show that the average of three observed interaction frequencies systematically underestimates the true regional mean interaction strength under the uniform-distribution model.

We apply the framework to avian frugivore–plant interaction data collected at three locations within each of six regions in Ecuador. The estimated distributions of interaction strength are used to test the hypothesis that species within the same genus spatially substitute for one another. The results do not support this hypothesis. Instead, species within the same genus tend to exhibit elevated interaction strengths in the same locations, indicating positive spatial association rather than substitution. We also find evidence that habitat fragmentation may reduce within-region variability in bird–plant interactions even when species diversity is maintained. These results suggest that explicitly modeling spatial variation can reveal ecological patterns that are obscured by conventional food-web representations.

## 1 Introduction

The food web is a fundamental concept in ecology. Interactions between consumer and consumed species are often summarized by measures of interaction strength, which are used to characterize community structure and ecosystem function. In practice, interaction strengths are commonly estimated from field observations collected at several sites within a regional habitat and then aggregated into a single value for each interacting pair of species. This procedure implicitly treats interaction strength as a fixed property of the region.

However, ecological interactions are influenced by local environmental conditions, species abundances, habitat structure, and other factors that may vary substantially within a region. As a result, interaction strengths observed at different sites often differ from one another. These differences arise partly from observational uncertainty and partly from genuine spatial variation in ecological interactions. Consequently, a regional interaction may be more appropriately viewed not as a fixed constant but as a spatially distributed latent variable whose value depends on location within the region.

Distinguishing between observational uncertainty and genuine spatial variation is important for understanding food-web structure. If interaction strengths vary substantially within a region, then a single aggregated estimate may obscure ecologically meaningful patterns. In such cases, the distribution of interaction strengths across locations becomes an important characteristic of the food web in its own right. Estimating this distribution requires methods that can separate measurement uncertainty from location-to-location variation.

In this paper, we develop a framework for estimating the probability distribution of food-web interaction strengths as a function of location within a given region. The framework uses site-specific observation data to estimate both variation among locations and uncertainty arising from the observation process. We assume a particular functional form for the distribution of location-dependent interaction strengths and estimate the parameters of that distribution together with their uncertainties.

The resulting estimates may be used to address a variety of ecological questions. One example concerns species that occupy similar ecological niches. Different species within the same genus may possess similar morphological features and feeding habits, potentially leading to functional overlap. If such species substitute for one another across locations, interactions involving those species should exhibit negative spatial correlations. Estimates of the spatial distribution of interaction strengths provide a means of testing this hypothesis and related questions concerning the organization of ecological communities.

More generally, our objective is to characterize regional food webs in terms of distributions of interaction strengths rather than single aggregated values. By treating interaction strength as a spatially distributed latent variable, we seek to quantify the extent of within-region heterogeneity and to determine what ecological information may be gained by explicitly accounting for spatial variation.

## 2 Background

Ecological interaction networks provide a quantitative framework for describing how species interact within communities and how these interactions influence ecosystem structure and function [Pimm, 1982, Cohen et al., 1990, Dunne et al., 2002]. Food webs in particular have become a central tool for understanding biodiversity maintenance, ecosystem resilience, and the effects of environmental disturbance on ecological communities [McCann, 2000, Pascual and Dunne, 2005]. Increasing attention has been devoted not only to the topology of ecological networks, but also to the strengths and variability of interactions among species [Berlow et al., 2004, Wootton, 1997].

In mutualistic and trophic systems alike, interaction strength is rarely spatially homogeneous. Local abundance, habitat structure, phenology, climate, and anthropogenic disturbance may all produce substantial variation in interaction frequencies across sites within the same region [Trøjelsgaard et al., 2015, Poisot et al., 2015]. Consequently, interaction networks constructed from pooled regional data may obscure important ecological heterogeneity occurring at smaller spatial scales [Olesen et al., 2011, Trøjelsgaard et al., 2015]. Spatial turnover in interactions may occur even when species composition remains relatively stable, indicating that ecological processes are influenced not only by species presence but also by local environmental context [Carstensen et al., 2014, Poisot et al., 2012].

Frugivore–plant interaction networks are especially sensitive to spatial variation because fruit availability and frugivore movement patterns fluctuate strongly across landscapes [Howe and Smallwood, 1982, Bleher and Böhning-Gaese, 2001]. In tropical regions, high species diversity and comparatively lower levels of network specialization can increase the number of potential interaction pathways, contributing to complex and spatially heterogeneous patterns of frugivory [Kissling et al., 2009, Schleuning et al., 2012]. Previous studies in Andean forests have shown that fragmentation and elevation gradients can alter both species composition and interaction structure [Quitián et al., 2018]. Such effects may influence not only the average interaction frequency between taxa but also the degree of variability among sites within the same regional habitat.

A major challenge in ecological network analysis is that observed interaction frequencies are subject to multiple sources of uncertainty [Martinez et al., 1999]. Sampling effort is necessarily limited, especially in species-rich tropical systems, and many interactions are rare or observed only intermittently [Blüthgen et al., 2008, Olesen et al., 2008]. Consequently, observed interaction matrices frequently contain large numbers of zeros that may reflect either genuinely absent interactions or incomplete sampling. Distinguishing observational uncertainty from true ecological variation therefore remains an important methodological challenge.

Several statistical approaches have been developed to address uncertainty in ecological interaction data. Bayesian methods have become increasingly important because they allow explicit representation of uncertainty and incorporation of hierarchical ecological processes [Ellison, 2004, Clark, 2005]. Bayesian frameworks are now widely used in population ecology, species-distribution modeling, and community ecology [Kéry and Schaub, 2011, McElreath, 2020], but comparatively fewer studies have focused on estimating uncertainty in food-web interaction strengths themselves.

Observed interaction frequencies are commonly modeled as count data using Poisson and related generalized linear models [McCullagh and Nelder, 1989]. These models arise naturally from assumptions about the occurrence of independent random events [McCullagh and Nelder, 1989]. However, ecological count data often exhibit overdispersion relative to Poisson expectations [Ver Hoef and Boveng, 2007]. One source of this heterogeneity is variation in species abundance, which has been shown to influence both interaction frequencies and interaction strengths in ecological networks [Vázquez et al., 2007]. More generally, ecological abundance distributions have frequently been described using lognormal and related statistical models [Preston, 1948, Sugihara, 1980].

Spatial heterogeneity in ecological interactions has also been studied through the concept of interaction turnover, or ‘β-diversity” of interactions [Poisot et al., 2012]. Interaction turnover may arise from species turnover, rewiring of interactions among co-occurring species, or environmentally mediated behavioral changes [Poisot et al., 2012, 2015, Caradonna et al., 2017]. Recent studies suggest that local environmental conditions can substantially reshape interaction networks even within relatively small geographic regions [Trøjelsgaard et al., 2015, Poisot et al., 2015]. These findings motivate the need for statistical methods capable of estimating not only average interaction strengths, but also the uncertainty and site-to-site variability associated with those strengths.

An additional question concerns the extent to which ecologically similar species substitute for one another across space. Functional redundancy has long been proposed as a mechanism contributing to ecosystem resilience [Walker, 1992]. Closely related species often share morphological and ecological traits due to phylogenetic niche conservatism, potentially allowing them to occupy similar ecological roles [Losos, 2008]. In frugivore systems, multiple bird species often interact with overlapping sets of fruiting plants, creating the potential for partial functional redundancy in seed dispersal [Bascompte and Jordano, 2006]. If such substitution occurs spatially, genus-level interaction strengths may exhibit lower site-to-site variation than would be expected from the summed variation of their constituent species-level interactions.

Despite growing interest in ecological network variability, comparatively few studies have attempted to estimate regional distributions of interaction strengths from sparse observational data. Most ecological network studies have instead focused on either aggregated networks or comparisons among site-specific networks [e.g., Bascompte and Jordano, 2006, Trøjelsgaard et al., 2015]. Methods that explicitly estimate latent regional interaction distributions and their associated uncertainties remain comparatively underdeveloped. The present study addresses this gap by introducing a Bayesian framework for estimating both uncertainty and intra-regional variation in bird–plant interaction strengths based on repeated observations from multiple locations within a region.

Our approach models observed interaction counts as Poisson-distributed measurements conditioned on latent interaction strengths that vary geographically according to an underlying probability distribution. By estimating the parameters of this latent distribution, we aim to distinguish measurement uncertainty from genuine ecological variability. We then apply this framework to frugivore–plant interaction data from multiple elevational and habitat contexts in southern Ecuador in order to investigate two related ecological questions: first, how strongly interaction strengths vary among sites within a region; and second, whether taxonomically related species tend to substitute for one another spatially.

## 3 Methods

In this section, we first present our modelling assumptions for the relation between interaction measurements and parameters that characterize the underlying interaction distributions within a given region. We then describe the general process of parameter estimation using a Bayesian framework. Next, we introduce the dataset to which we apply the model. This data includes transect measurements taken from six regions in Ecuador. Finally, we give the particular form of the estimation equations that can be used for this dataset. In particular, we consider two possible forms of the geographical interaction rate distribution (Poisson and uniform), and argue that the uniform model is superior, based on preliminary observations.

### 3.1 Assumptions for the mathematical model

We suppose an ecologically homogeneous region in which transect survey data for bird-fruit interaction events is taken at *M* specific locations. We denote the *M* observed interaction counts as the vector *c⃗* := [*c*_1_, …, *c_M_*]. The bird-fruit interaction counts are assumed to be Poisson distributed, where the Poisson rate depends on the actual interaction strength. Specifically, if the actual strength at location *m*(1 ≤ *m* ≤ *M*) is λ*_m_*, then the probability distribution of interaction counts is given by:

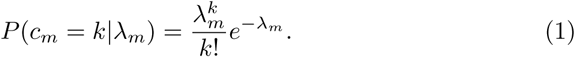

The actual interaction strengths {λ*_m_*} differ among the *M* locations according to an underlying distribution which gives the probability density for the interaction strength at a randomly-chosen point within the region. The *M* interaction strengths λ_1_ … λ*_M_* are assumed to be statistically independent. The underlying distribution will have a specific form, and depend on a set of parameters. We notate this underlying distribution by p(λ|*q⃗*), where *q⃗* = [*q*_1_, …*q_N_*] are *N* parameters that characterize the underlying distribution: for example, if the underlying distribution is assumed to be normal, then the mean and standard deviation would be parameters that uniquely specify the distribution.

### 3.2 Bayesian estimation of location-dependent food web interaction strengths

The task at hand is to estimate the parameter vector *q⃗* based on the measurement vector *c⃗*. We use a standard Bayesian approach as follows. Bayes’ formula applied to the vectors *c⃗* and *q⃗* gives:

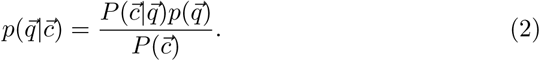

In (2), the small ‘*p*’ quantities represent probability densities, while the large ‘*P*’ quantities represent probability values in a discrete distribution (recall *q⃗* is continuous, but *c⃗* is discrete). It follows from (2) that the conditional probability distribution of the parameter vector *q⃗* given the set of measurements *c⃗* can be computed based on the conditional probability distribution *P*(*c⃗*|*q⃗*) and the prior probability density *p*(*q⃗*) and prior distribution *P*(*c⃗*). *P*(*c⃗*|*q⃗*) can be computed based on (1) and the assumption that {λ*_m_*} are independent and identically distributed as:

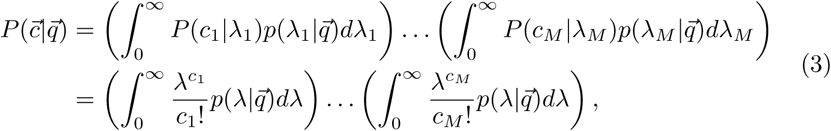

where the subscripts are removed from the λ variables in the second equation because they are dummy variables that are integrated out.

To complete the evaluation of *p*(*q⃗*|*c⃗*), it remains to find *P*(*c⃗*) and *p*(*q⃗*), which are the prior probabilities for *c⃗* and *q⃗*. Since we have no prior knowledge pertaining to *c⃗* and *q⃗*, we assume they have uniform distributions. It follows that the ratio *p*(*q⃗*)/*P* (*c⃗*) is taken as a constant. This gives:

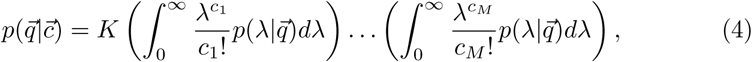

where the constant *K* is determined by normalization conditions,

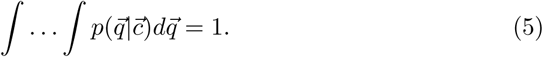

We have not yet assigned a specific form to *p*(λ|*q⃗*), which appears in (4). In the next section, we introduce a specific dataset that will be analyzed using the above procedure and discuss an appropriate choice of *p*(λ|*q⃗*) for this dataset.

### 3.3 Dataset

Data was collected from 18 1-hectare plots spread across three elevations (1000 m above sea level (asl) (970–1092 ± 47.2 m), 4*^◦^* 6*^′^* S, 78*^◦^* 58*^′^* W; 2000 m asl (2025–2119 ± 32.4 m), 3*^◦^* 58*^′^* S, 79*^◦^* 4*^′^* W; and 3000 m asl (2874–2966 ± 32.9 m], 4*^◦^* 6*^′^* S, 79*^◦^* 10*^′^* W. These plots represented two types of habitat (natural forest and fragmented forest), within and around Podocarpus National Park (PNP) and San Francisco reserve (BRSF) on the southeastern slope of the Andes in Ecuador. The six sites are characterized in Table 1. Study plots were selected jointly by the German Platform for Biodiversity and Ecosystem Monitoring and Research in South Ecuador, to ensure typical representation of each forest type. The area exhibits a humid tropical montane climate [Kottek et al., 2006] with a bimodal rainfall pattern (highest precipitation season from May through June, lowest precipitation season from October through November) [Emck, 2007]. The mean annual temperature is 20 *^◦^*C and the mean annual precipitation is 2432 mm at low elevations. At mid elevations, the mean annual temperature is 15.5 *^◦^*C, and the mean annual precipitation is 2079 mm. At high elevations, the mean annual temperature is 10 *^◦^*C, and the mean annual precipitation is 4522 mm [Emck, 2007]. The study plots encompass three distinct vegetation types: evergreen premontane forest at low elevations, evergreen lower montane forest at mid elevations, and upper montane forest at high elevations [Homeier et al., 2008]. The vegetation composition varies across elevations, with emergent trees, climber plants, and lianas dominating at low elevations, a few emergent trees, vascular epiphytes, and climber plants at mid elevations, and a high percentage of Weinmannia shrubs, vascular epiphytes, and epiphytic mosses at high elevations [Paulsch et al., 2008]. The dataset is described in more detail in reference Thron et al. [2023].

**Table 1:** Summary of characteristics for six regional habitats studied.

| Location | Approx. altitude (m) | Environment type |
| --- | --- | --- |
| Bombuscaro | 1000 | Natural |
| Copalinga | 1000 | Fragmented |
| ECSF | 2000 | Natural |
| Finca | 2000 | Fragmented |
| Cajanuma | 3000 | Natural |
| Bellavista | 3000 | Fragmented |

Figure 2 (see Appendix) shows histograms for bird-fruit interactions by region. Figure 2a shows the maximum interaction strength (*c_max_* := max(*c*_1_, *c*_2_, *c*_3_)) between bird and fruit species among the three locations in each region, while Figure 2b shows the maximum interaction strength between bird and fruit genera. The vertical scale for Figure 2b is smaller than that for Figure 2a, reflecting the fact that there are fewer genera than species. Since *c_max_* varies over three orders of magnitude, the horizontal axis of the histogram is given in logarithmic scale, and the histogram bins are also logarithmic.

The histogram shapes are roughly similar between regions, with large counts for the lowest bins (≤ 3 interactions) and subsequently decreasing steadily as a function of the number of interactions. The histograms are also roughly similar between species- and genus-level analyses, except as expected, the genus histograms show a larger proportion of counts in the higher bins.

To analyze the data from this dataset using our Bayesian approach, it is necessary to specify a form for *p*(λ|*q⃗*). In the following subsection, we discuss two alternative possibilities and rule out one of the alternatives.

### 3.4 Alternative forms for interaction strength distribution

The interaction strength distribution *p*(λ|*q⃗*) is a key quantity in the estimation equation (4). However, neither λ nor *q⃗* can be accessed directly. Therefore we must rely on general considerations to inform our choice of *p*(λ⃗|*q⃗*). In this section, we consider some alternative possibilities and evaluate their suitability.

#### 3.4.1 Poisson distribution

Our first choice of distribution relies on the assumption that the interactions between each bird–plant species pair within a region are uniformly distributed geographically. In this case, the total number of interactions per time in each location would correspond to the interactions within a subarea of the region of given size. This situation is depicted in Figure 1 where the interaction rate λ would be Poisson distributed over the set of locations within the region.

**Figure 1:**
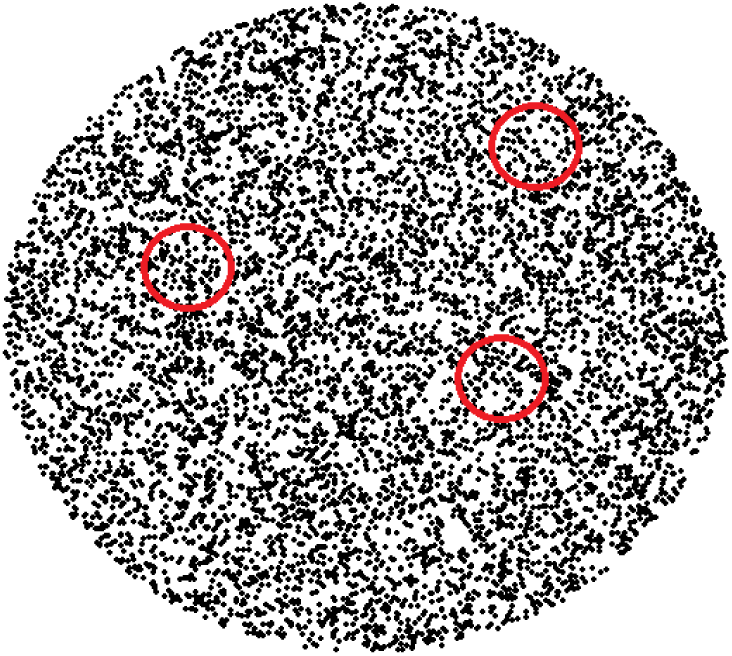
Schematic illustration of Poisson model for geographical variation of interaction strengths. The black dots represent the bird-plant interactions within a circular region. Each red circle represents a site within the region where measurements are taken, and the dots enclosed in the circle represent the number of actual interactions at that site. (Measured interactions would comprise a small fraction of the total number of actual interactions.)

One consequence of this model is that variations in interaction strength between sites are relatively small. For example, a Poisson random variable with a mean rate of 14 has a less than one in a million chance of attaining a value of 0. However, in the dataset we find many instances of species that have interactions strengths of 20 or more in one location within a region, and zero at another location within the same region (see e.g. Figure 10). Such occurrences are extremely unlikely under the assumption of Poisson distribution, so we reject the Poisson distribution as a possible form for p(λ|q⃗). In the next subsection, we will present an alternative that is much more reasonable.

#### 3.4.2 Normal and lognormal distributions

A natural alternative to the Poisson model is to assume that interaction strengths are distributed according to either a normal or a lognormal distribution across locations within a region. Such distributions are commonly used in ecological applications because they allow substantially greater variation than the Poisson distribution and can accommodate heterogeneous environments.

The normal distribution, however, has an important conceptual drawback for interaction strengths. Since interaction rates cannot be negative, a normal distribution necessarily assigns positive probability to biologically impossible values unless the variance is sufficiently small relative to the mean. Restricting the variance to avoid this problem would again limit the model’s ability to explain the large differences in interaction strength that are frequently observed between nearby sampling locations.

The lognormal distribution avoids negative values and is often used to model ecological quantities that vary over several orders of magnitude. However, its strongly right-skewed form implies a long upper tail. Consequently, the distribution assigns non-negligible probability to interaction strengths that are much larger than those observed unless the variance parameter is tightly constrained. In practice, this introduces an additional parameter that must be estimated reliably. Since our data consist of observations from only three locations for each bird–plant interaction, estimation of both the location and scale parameters of a lognormal distribution is expected to be unstable and highly sensitive to sampling variation.

A further consideration is mathematical complexity. The Bayesian estimation procedure developed in Section 3.2 requires repeated evaluation of integrals of the form appearing in (4). For a uniform interval distribution these integrals can be evaluated efficiently using incomplete gamma functions. Under either the normal or lognormal model, the corresponding integrals involve products of Poisson likelihoods with Gaussian or lognormal densities and do not simplify to comparably convenient expressions. The resulting numerical integration is considerably more computationally demanding while providing little additional information given the limited amount of data available for each interaction.

For these reasons, although normal and lognormal distributions are plausible candidates in settings where substantially more observations are available, they appear less suitable for the present study.

#### 3.4.3 Uniform interval distribution

We have noted that measured interaction rates vary widely from location to location in the same region. We expect therefore that the interaction rate distribution will also be wide, and not narrowly peaked. One common example of a wide distribution is the uniform distribution on an interval. Some reasons for using a uniform distribution are given as follows.

First, without detailed information about the region there is no particular reason to suppose that there is one interaction rate that is more likely than another (within reasonable bounds). It is true that within ecology, the use of uniform priors is discouraged, but this is because a uniform prior with a large upper bound allows for impossibly high interaction rates [Lemoine, 2019]. In our case, we can limit the range of interaction rates by specifying an upper and lower bound to the uniform distribution, which will be estimated by the measurements.

Second, the uniform interval distribution is specified by only two parameters. In our case, having only three locations means that we only have three data points that can be used to estimate the distribution parameters for each interaction. So, it is not possible to estimate a distribution with too many parameters.

Third, the implementation of a uniform interval distribution estimator is comparatively simple. Let q_1_ and q_2_ be the lower and upper endpoints, respectively, of the interval that we are estimating. We may specify a prior range for the values as

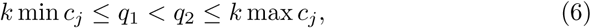

where *k* is a constant greater than 1 (in our implementation, we used *k* = 2). We will also assume a maximally uninformative prior distribution for *q⃗* := [*q*_1_, *q*_2_] in this range. So we have:

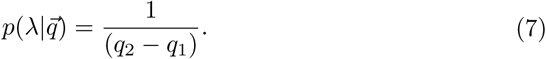

Substituting (7) into (4) with *M* = 3 gives

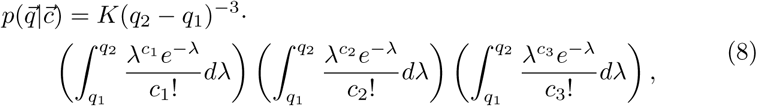

where *c⃗* := [*c*_1_, *c*_2_, *c*_3_]. We may rewrite this as:

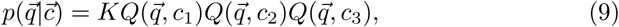

where

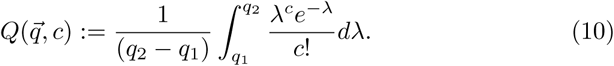

Note that *Q*(*q⃗*, *c*) is the average value of 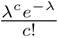 on the interval [*q*1, *q*2]. It may be verified (see Appendix) that

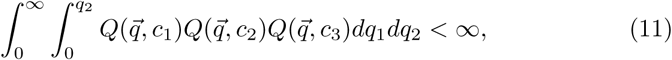

so it is possible to find a value of *K* > 0 that will normalize the conditional distribution *p*(*q⃗*|*c⃗*) so that it has total mass 1:

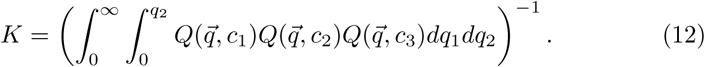

#### 3.4.4 Conclusions

In view of above discussions, the uniform interval distribution appears to represent a practical compromise between biological realism and statistical tractability. Unlike the Poisson model, it permits large differences in interaction strength between locations. Unlike the normal distribution, it assigns no probability to negative interaction rates, and unlike the lognormal distribution, it does not impose a long upper tail that must be estimated from very limited data. Moreover, because it is described by only two parameters, it can be estimated more reliably from observations at only three locations than models requiring additional shape parameters. For these reasons, we adopt the uniform interval distribution as the working model for the remainder of this study.

### 3.5 Heatmap representations for distribution characteristics

Based on the findings described in Section 3.4, the following analysis assumes a uniform interval distribution as the underlying form for the variation of species-to-species interaction strength within each region. It follows that (8) governs the estimation of the underlying parameter vector *q⃗* from field observations.

To provide further insight into the behavior of the distribution (8), we construct several heatmaps that show the dependence of statistical parameters as a function of the vector of observations *c⃗*. The x- and y-axes of each heatmap represents the ratios *c*_1_/*c*_2_ *c*_2_/*c*_3_ respectively, where *c*_1_, *c*_2_, *c*_3_ are the three interaction strength measurements and *c*_1_ ≤ *c*_2_ ≤ *c*_3_. Since 0 ≤ *c*_1_/*c*_2_, *c*_2_/*c*_3_ ≤ 1, it follows that the x and y limits on the heatmaps are both [0,1]. Note that *c*_3_ > 0 so *c*_2_/*c*_3_ is always defined, and when *c*_1_ = *c*_2_ = 0, we set *c*_1_/*c*_2_ equal to a random value between 0 and 1, to avoid data points stacking on top of each other.

Since the magnitude (and to some extent the shape) of the heatmap contours depends also on the value of the largest measurement c_3_, we show plots for four different values of *c*_3_ (15,25,35,45). Each heatmap shows the dependence on *c*_1_, *c*_2_, *c*_3_ of a different statistical characteristic of the inferred probability distributions. The characteristics chosen reflect bias, precision, and variability of estimators derived from the distribution. These characteristics include:

a. Estimated mean interaction strength for the region, divided by the mean of observed interaction strengths. Mathematically, this is given by:

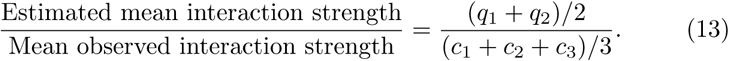

This ratio shows whether or not the mean of observed interactions is an unbiased estimator for the actual mean interaction strength in the region, given the assumption of an underlying uniform interval distribution.
b. 90% confidence interval width for the mean interaction strength, divided by the estimated mean interaction strength. This quantity indicates the precision of the estimate of mean regional interaction strength, based on measurements in three locations.
c. Estimated range of location-specific interaction strength, divided by the estimated mean interaction strength for the region. This ratio quantifies the intra-regional variability of interaction strength from location to location.
d. Uncertainty in the estimated range of location-specific interaction strength, divided by the estimated mean interaction strength of the region. This ratio quantifies the relative uncertainty in the intra-regional variability estimated in (c).

For each bird-fruit interaction in the six different regions, we compute the ratios *c*_1_/*c*_2_ and *c*_2_/*c*_3_ and display the point (*c*_1_/*c*_2_, *c*_2_/*c*_3_) on the appropriate heatmaps. Since the estimation of *p*(*q⃗*|*c⃗*) depends only on c⃗ and not on specific regional characteristics, we are justified in displaying (*c*_1_/*c*_2_, *c*_2_/*c*_3_) values for all interactions for all regions on the same plots. Each bird-fruit interaction is plotted on only one of these graphs, depending on the value of *c*_3_ (equal to *c_max_*) for the interaction. For the ranges 10 ≤ *c*_3_ ≤ 20, 20 ≤ *c*_3_ ≤ 30, 30 ≤ *c*_3_ ≤ 40, and *c*_3_ ≥ 40, we plot the interaction first, second, third or fourth plot respectively.

### 3.6 Species substitution by genus

We have mentioned that some of the observed variation in interaction strength from site to site within a region may be due to measurement error, and some may be due to real differences in interaction strength. It is possible that the differences in species-species interaction strength within a regional habitat occur because different species with similar functional characteristics fill the same ecological niche in different locations within the region. For example, suppose bird species *A*_1_ and *A*_2_ and plant species *B*_1_ and *B*_2_ are functionally similar. It is possible for example that at one location, *A*_1_ interacts strongly with *B*_1_ and *B*_2_ and *A*_2_ has a weak presence, while at another location *A*_2_ interacts strongly with the same species, and *A*_1_ shows weak or nonexistent interaction. In other words, species *A*_1_ and *A*_2_ effectively substitute for each other in the two respective regions.

Using our model, we may quantify the extent to which species substitution occurs within bird and plant genera. If species substitution occurs, then the variance of genus-level interaction strength among locations should be less than the sum of individual variances within the genera. We may calculate these variances for several bird-plant genera pairs, and see if there is a general tendency in the relative sizes of the per-genus variance versus the sum of included per-species variances.

## 4 Results

### 4.1 Heatmaps of location-dependent distribution characteristics

In this subsection, we analyze the heatmap representations described in Section 3.5. All figures discussed are contained in the Appendix.

#### Midpoint distribution

Figure 3 gives heatmaps for the ratio of the estimated midpoint of the interaction strength to the mean of the three observed interaction strengths. In each graph, the four corners (0, 0), (0, 1), (1, 0), and (1, 1) correspond to *c*_1_ ≪ *c*_2_ ≪ *c*_3_, *c*_1_ = *c*_2_ ≪ *c*_3_, *c*_1_ ≪ *c*_2_ = *c*_3_, and *c*_1_ = *c*_2_ = *c*_3_ respectively. The points in each plot represent observed (*c*_1_*, c*_2_*, c*_3_) values in the different regions. For example, in Figure 3a the black square near (0.5, 0.4) in the first panel indicates that there was a species-species interaction in Copalinga for which 10 ≤ *c_max_* ≤ 20 and the three measurements *c*_1_ ≤ *c*_2_ ≤ *c*_3_ satisfied the relations *c*_1_ ≈ 0.5*c*_2_ and *c*_2_ ≈ 0.4*c*_3_ (thus *c*_1_ ≈ 0.2*c*_3_). As another example, the red circle located near (0.2, 0.6) in the fourth panel of Figure 3b indicates there was a bird genus-fruit genus interaction in ECSF with maximum interaction strength *c_max_* greater than 40, for which the three measurements *c*_1_ ≤ *c*_2_ ≤ *c*_3_ satisfied the relations *c*_1_ ≈ 0.2*c*_2_ and *c*_2_ ≈ 0.6*c*_3_ (hence *c*_1_ ≈ 0.12*c*_3_). In all figures, a point on the lower edge with *c*_2_/*c*_3_ = 0 indicates that the interaction strength was 0 at 2 locations within the region (the points with *c*_1_ = *c*_2_ = 0 are plotted with random values of *c*_1_/*c*_2_ in order to distinguish them).

A comparison of the panels in Figures 3 shows that the heatmap contours are relatively independent of the magnitude of *c_max_*. This implies that the ratio of estimated mean interaction strength to mean of observed interaction strengths depends mostly on the relative sizes of the measured interaction strengths, rather than the absolute sizes.

Note that the expected-midpoint-to-observation-average ratio is consistently greater than 1, and only approaches 1 when *c*_2_/*c*_3_ is close to 1. This means that the mean of observed interaction strengths tends to underestimate the actual interaction strength in the region, particularly if *c*_3_ ≫ *c*_2_ (i.e. *c*_2_/*c*_3_ is close to 0). In the limit as *c*_2_/*c*_3_ → 0, then the estimated mean of the distribution interval is roughly twice the average of the three measurements, regardless of the value of *c*_1_.

When comparing the results for species and genus (Figures 3a and 3b, respectively), we note that there are more interactions in the lowest bin (10 ≤ *c_max_* ≤ 20) for species compared to genus, and there are more interactions in the highest bin (*c_max_* ≥ 40) for genus compared to species. This is to be expected, because a genus consists of several species so we expect larger interaction counts for genus than species.

In both the species and genus *c_max_* > 40 panels we find points with *C*_2_/*C*_3_ = 0. This indicates species and genus pairs for which no interactions are observed at two locations, and more than 40 are recorded at a third location. This demonstrates the possibility of wide variations in interaction strength from location to location within a region, as noted in Section 3.4.1.

#### Midpoint uncertainty

Figure 4 shows the width of 90% confidence intervals for the mean interaction strength, as a function of interaction strength ratios *c*_1_/*c*_2_ and *c*_2_/*c*_3_. The width is shown as a proportion of the estimated mean interaction strength.

#### Location-specific interaction strength standard deviation

The model we have proposed takes into account that interaction intensities vary from place to place in a single environment. The dependence of the standard deviation of the location-dependent interaction intensity on the three location measurements is shown in Figure 5. The figure shows the ratio of estimated standard deviation values to the estimated mean interaction strength value. If both *c*_2_/*c*_3_ and *c*_1_/*c*_2_ are small, the ratio can range up to 2, meaning there is a very large predicted variation in interaction strength across the environment. Even when the measurements are close to each other, the standard deviation is still considerable, about 40% of the estimated mean.

#### Uncertainty in location-specific interaction strength distribution width

The estimated standard deviation depicted in Figure 5 has a large uncertainty. Figure 6 shows this uncertainty as a proportion of the estimated mean interaction strength for different measurement ratios. The large uncertainties show that standard deviations for location-dependent intensity distributions cannot be reliably estimated based on measurements in three locations.

### 4.2 Bar plots

The panels in Figure 7 summarize uncertainties in frugivore bird species-fruit species interaction strength by region. In each region, frugivore bird species-fruit species pairs are listed in order of increasing interaction strength. For each region, we exclude species-species interactions whose maximum observation count among the three locations is less than 10. Each bar represents one species-species interaction. The horizontal black segment marks the estimated mean strength of the given interaction in the given region. The blue bar shows the 90% confidence interval for the mean strength of interaction. The red bar shows the 90% confidence interval for location-specific interaction strength within the given region. Figure 8 similarly represents uncertainty information for genus-genus interactions in the six different regions.

Comparison of the top two panels of Figures 7 and 8 shows that there is less variability (and slightly lower interaction levels) for species and genera in Copalinga compared to Bombuscaro. This indicates that there is less site-to-site variability in Copalinga (which is at the same altitude as Bombuscaro but whose habitat is fragmented by human activity).

### 4.3 Species substitution plots

Figure 9 tests the hypothesis of species substitution, by plotting both the variance of genus-genus interaction strengths (orange lines) and the sums of variances for species-species interactions for species included within the corresponding genus (blue lines). Species are omitted if the total number of interactions is less than 10. Data is plotted for genus pairs within four of the six regions: insufficient data was present for Cajanuma and Bellavista. The results clearly show that the variance per genus consistently exceeds the sum of variances of constituent species, contradicting the hypothesis that species within genera substitute for each other at different locations.

Figure 10 reinforces this conclusion by displaying the species-species interaction strengths as stacked bars, where the bars correspond to different genera. Each pair of (frugivore, fruit-bearing plant) genera have three stacked bars, corresponding to the three observation locations. For several genus pairs, we find a single dominant, multicolored bar, indicating one of the three locations within a region for which several species congregate while the other two locations have few or no recorded interactions representing the same genus pair. This is particularly striking in the [Tangara, Indet], [Chlorospingus, Miconia], and [Tangera, Miconia] genus pairs in Bombuscaro. Partial species substitution may be observed in some cases, notably in the [Tangara, Miconia] genus pair in ECSF: in this case, there are two locations in which two completely different sets of species predominate. However, the species substitution is limited to two locations, and the third location has low interaction strength for all species within the genera.

Figure 10 also indicates species diversity profiles for each site and each bird genus-plant genus combination as a pair of numbers (a, b) above each bar, where a and b represent the number of bird and plant species respectively that were observed at the site from the given genus-genus combination. Diversity profiles within genera varied slightly between different Bombuscaro sites: for example, Tangara had 6, 6, and 4 species at the three different sites, while Miconia had 4, 3, and 2 species at the same three sites. On the other hand, interaction levels for these genera varied widely. At the most active site (which also showed the greatest diversity in both bird and plant species), over 500 interactions were recorded, while the least active site (which was also the least diverse) had fewer than 50 interactions.

Compared to Bombuscaro, Copalinga had more uniform levels of interaction among the three sites, with comparable species diversity. Diversity profiles at the different Copalinga sites are also quite similar: the site showing most interactions had increased interactions across all species compared to other sites.

The two mid-altitude sites (ECSF and Finca) had considerably less activity than the low-altitude sites, and no site recorded more than 200 interactions. At ECSF, there were two sites that appear to show species substitution within (tangara, miconia) interactions.

## 5 Discussion

The primary objective of this study was to develop a probabilistic framework for estimating uncertainty in bird–plant interaction strengths when measurements are available from only a small number of locations within a region. The results demonstrate that, under a Bayesian framework, it is possible to estimate not only mean interaction strengths but also the uncertainty associated with spatial variation in those interaction strengths. Rather than treating observed interaction counts as exact measures of regional interaction intensity, the proposed approach explicitly models the uncertainty arising from geographical heterogeneity.

The analysis also demonstrates that this uncertainty is substantial. Confidence intervals for mean interaction strengths are generally wide, while the uncertainty associated with location-specific interaction strengths is larger still. These findings suggest that interaction strengths estimated from only a few sampling sites should be interpreted with caution, particularly when they are used in quantitative analyses of ecological interaction networks. An additional consequence of the model is that the average of the observed location counts tends to underestimate the true regional interaction strength, especially when one location exhibits substantially higher interaction rates than the others. This suggests that simple averaging of observed counts may systematically underestimate regional interaction strengths whenever appreciable spatial heterogeneity is present.

The present study has several important limitations. Most notably, only three sampling sites were available within each region, which is one of the principal causes of the large uncertainties in the estimated interaction strengths. In addition, the analysis is based on a single probability model for the geographical distribution of interaction strengths. Although we argued that a uniform interval distribution provides a reasonable compromise between biological realism and statistical tractability for the present dataset, this assumption has not yet been validated using independent field observations. Consequently, the quantitative conclusions presented here should be regarded as tentative. Nevertheless, the Bayesian estimation framework itself is sufficiently general that alternative probability models can readily be substituted and compared using the same methodology.

Despite these limitations, the present work demonstrates that uncertainty arising from spatial variation in interaction strength can itself be estimated. Although the resulting confidence intervals remain wide with the present dataset, explicitly quantifying this uncertainty represents an important step beyond treating observed interaction counts as exact measures of regional interaction strength. More generally, the proposed framework provides a means of incorporating uncertainty directly into quantitative analyses of ecological interaction networks.

There are several promising directions for future work. First, the robustness of the present conclusions should be investigated with respect to the assumptions of the statistical model. In particular, it will be important to determine whether the estimated interaction strengths and associated confidence intervals remain stable when alternative distributions, such as lognormal, gamma, or mixture distributions, are used in place of the uniform interval distribution adopted here. Such analyses would help distinguish conclusions that are intrinsic to the data from those that depend on the modeling assumptions.

Second, validation of the proposed framework will require substantially larger datasets containing measurements from many more locations within each region. Such datasets would permit the use of separate training and testing sets for parameter estimation and model validation, allowing direct assessment of predictive accuracy. They would also make it possible to compare competing distribution models empirically rather than relying primarily on theoretical considerations. Finally, larger studies would allow investigation of how uncertainty in interaction strength varies with sampling effort and spatial scale, thereby providing practical guidance for the design of future ecological interaction surveys. Ultimately, improved probability models together with larger and more spatially extensive datasets should permit increasingly accurate estimation of interaction strengths and their uncertainties, leading to more reliable quantitative descriptions of ecological interaction networks.

## Data Availability

The datasets generated and/or analyzed during the current study are not publicly available but are available from the corresponding data custodian upon reasonable request. Requests for access to the data should be directed to Vinicio Santillán.

## Code Availability

The computer code used to implement the methods described in this study and to reproduce the simulation results is publicly available at https://github.com/AaronTAMUCT/Uncertainty.

## Use of Generative AI and AI-assisted Technologies

During the preparation of this manuscript, the authors used ChatGPT (OpenAI) to assist with literature discovery, critical evaluation of the manuscript, improvement of the organization and clarity of the writing, suggestions for alternative wording, and identification of grammatical issues. The AI system was not used to generate data, perform statistical analyses, derive mathematical results, or formulate the scientific conclusions. References suggested by the AI system were independently evaluated by the authors before inclusion. All AI-generated suggestions were critically reviewed, revised as appropriate, and verified by the authors. The authors take full responsibility for the content of the published manuscript.

## CRediT statement

**Conceptualization**: V.S.,A.B.,C.T.; **Data curation**: V.S., M.Q., E.L.N., M.S.; **Formal analysis**: A.B.,C.T.; **Funding acquisition**: K.B.S.,O.A.; **Investigation**: V.S., M.Q., E.L.N., M.S.; **Methodology**: A.B.,C.T.; **Project administration**: V.S., K.B.S.,O.A.; **Resources**: K.B.S.,O.A.; **Software**: A.B.,C.T.; **Supervision**: V.S., C.T.; **Validation**: A.B.,C.T.; **Visualization**: A.B.,C.T.; **Writing – original draft**: A.B.,C.T.; **Writing – review & editing**: A.B.,V.S., K.B.S., C.T.

## Appendix: Proof of integral convergence

In this appendix, we prove the following generalization of (11):

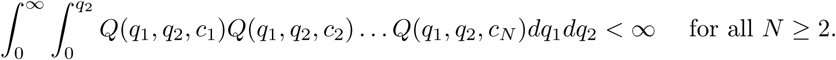

This inequality may be proved as follows. Denoting the integrand in (11) as *Q*, we have:

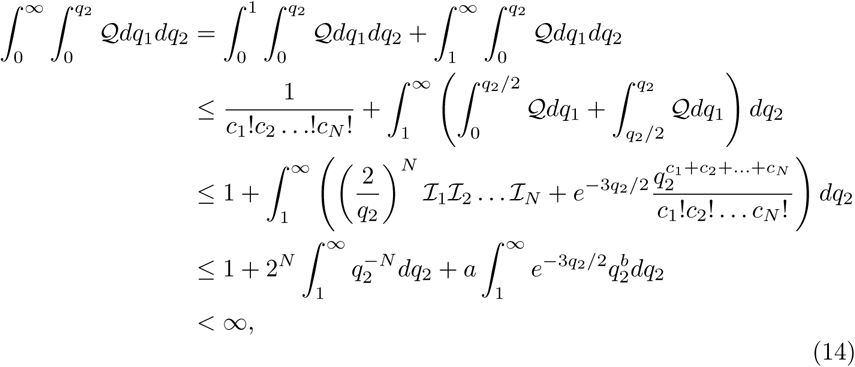

Where 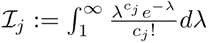 and *a*, *b* are nonnegative constants.

It is noteworthy that the inequality cannot be established for *N* = 1. Thus a single measurement is not sufficient to estimate the distribution parameters *q*_1_, *q*_2_ without an informative prior.

### Appendix: Figures

**Figure 2:**
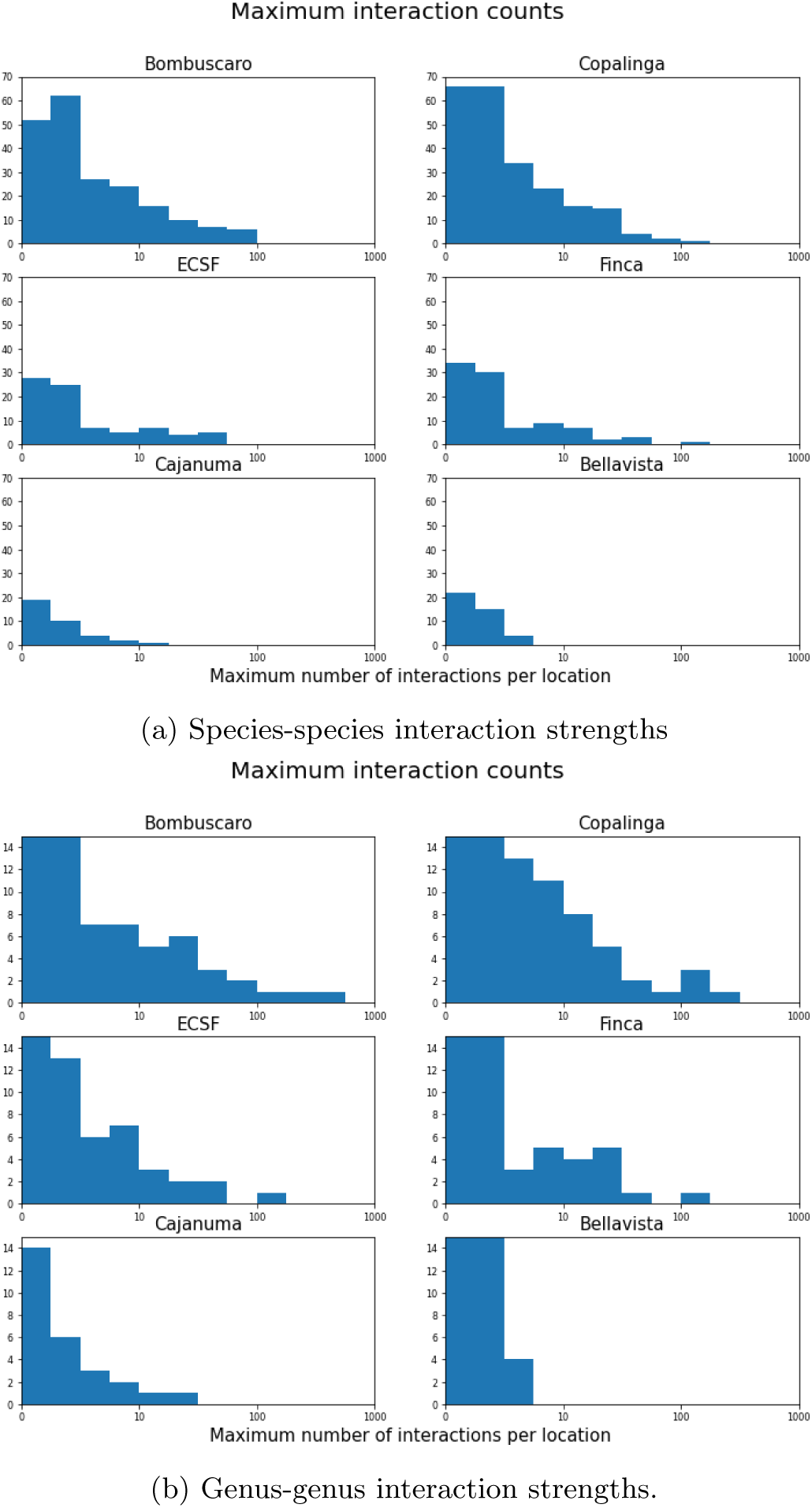
Histograms of interaction strengths by region. Species-species interaction strengths are shown in Subfigure (a), while genus-genus interactions are shown in Subfigure (b).

**Figure 3:**
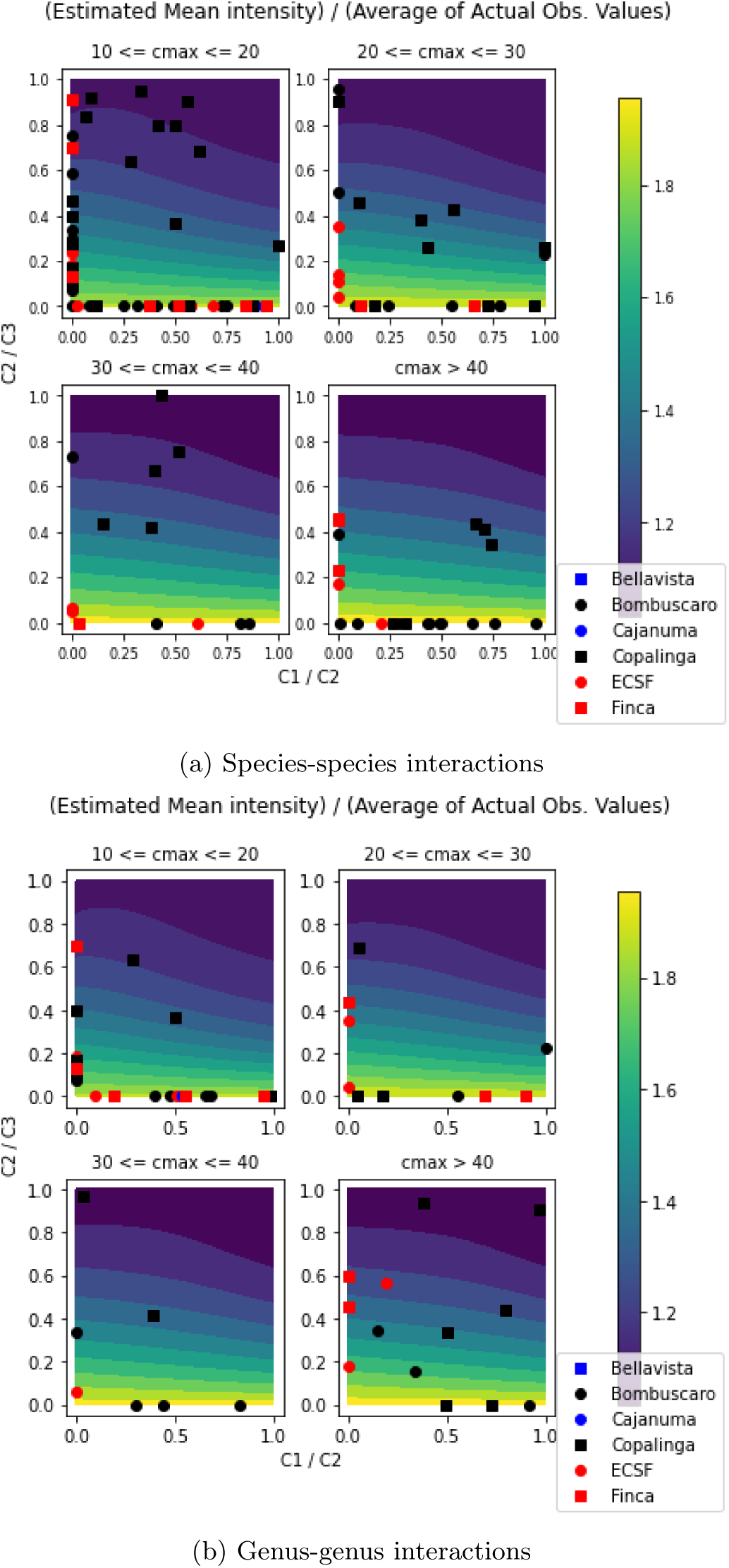
Ratio of estimated distribution mean to mean of three measurements as a function of interaction strength ratios *c*_1_/*c*_2_ and *c*_2_/*c*_3_. The plot also indicates interaction strength ratios for different species and different genera, grouped according to maximum interaction strength *c_max_*.

**Figure 4:**
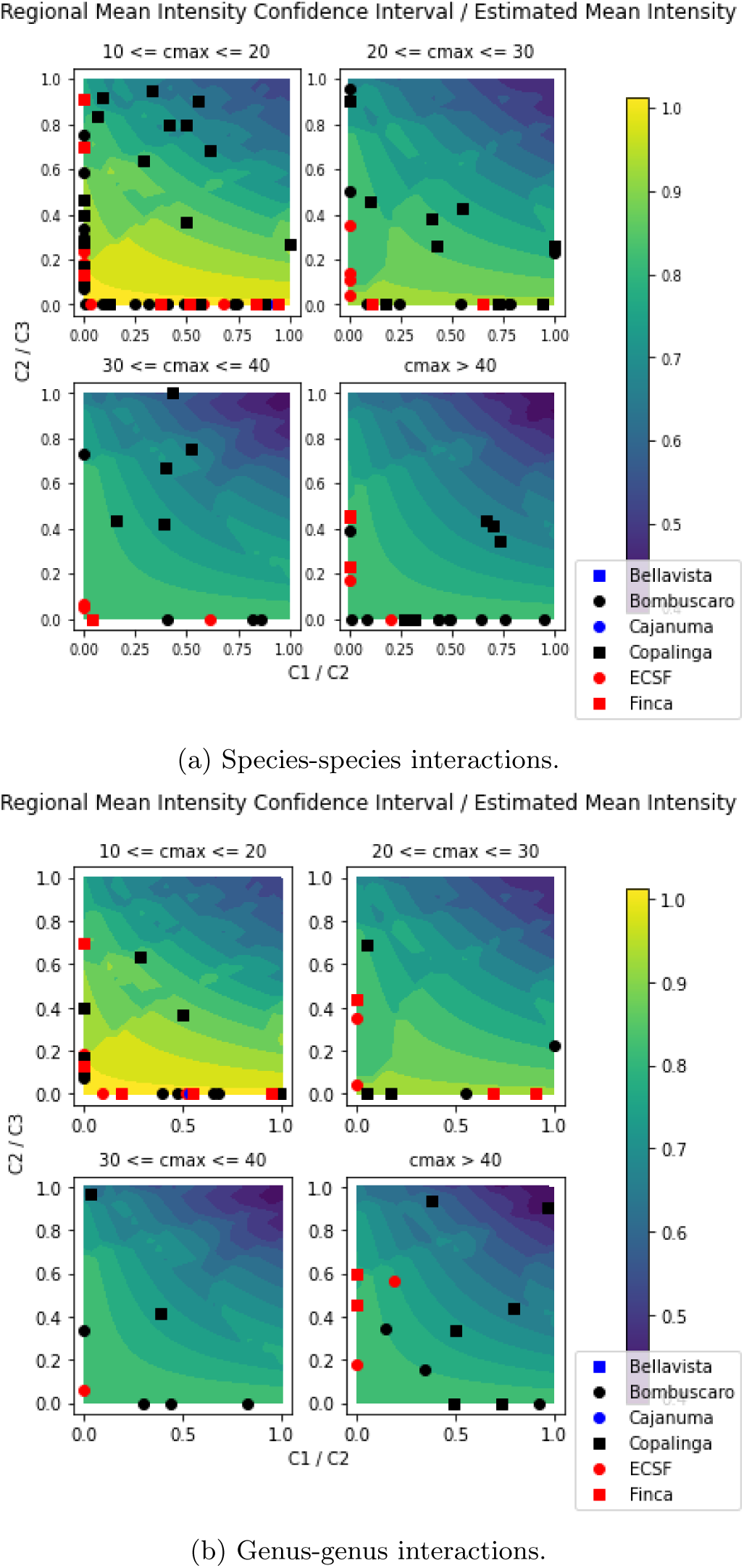
Ratio of 90% confidence interval for mean interaction strength to estimated mean interaction strength as a function of interaction strength ratios *c*_1_/*c*_2_ and *c*_2_/*c*_3_. The plot also indicates interaction strength ratios for different species and different genera, grouped according to maximum interaction strength *c_max_*.

**Figure 5:**
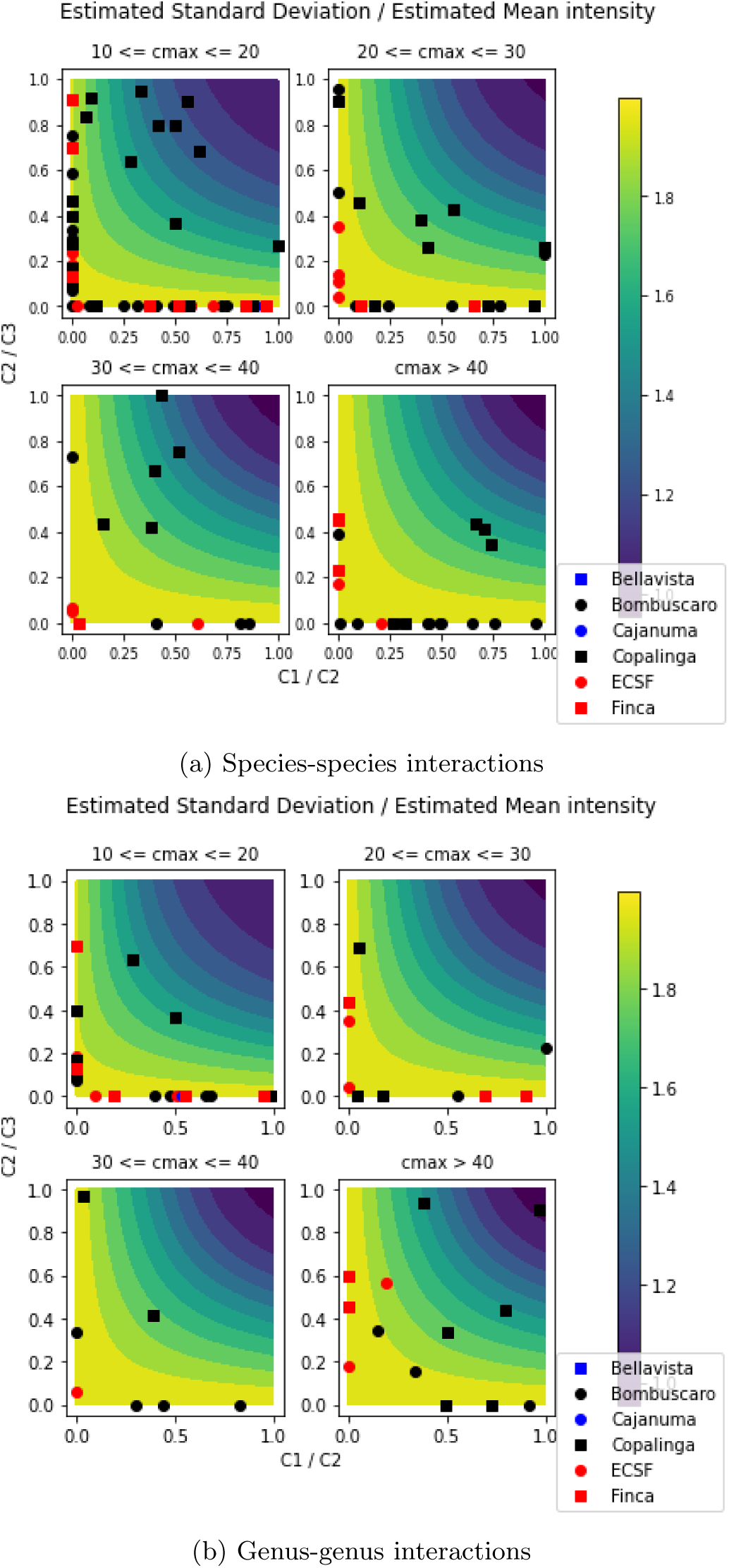
Estimated ratio of range of standard deviation of location-specific intensities to estimated mean interaction strength, as a function of observed interaction strength ratios *c*_1_/*c*_2_ and *c*_2_/*c*_3_. The points on the plot also indicate interaction strength ratios for different species and different genera, grouped according to maximum interaction strength *c_max_*.

**Figure 6:**
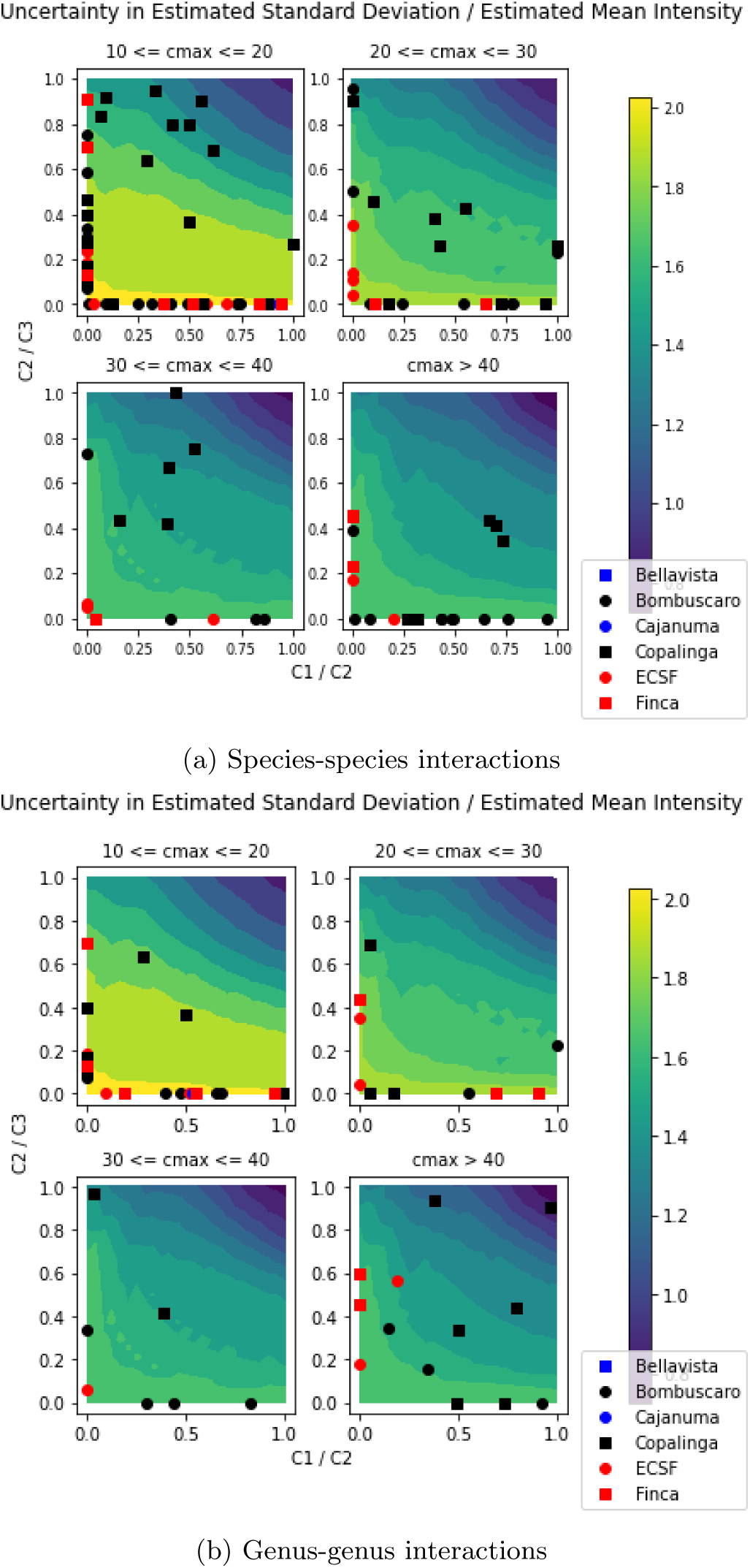
Ratio of 90% confidence interval for standard deviation to estimated mean interaction strength, as a function of observed interaction strength ratios *c*_1_/*c*_2_ and *c*_2_/*c*_3_. The points on the plots also indicates interaction strength ratios for different species and different genera, grouped according to maximum interaction strength *c_max_*.

**Figure 7:**
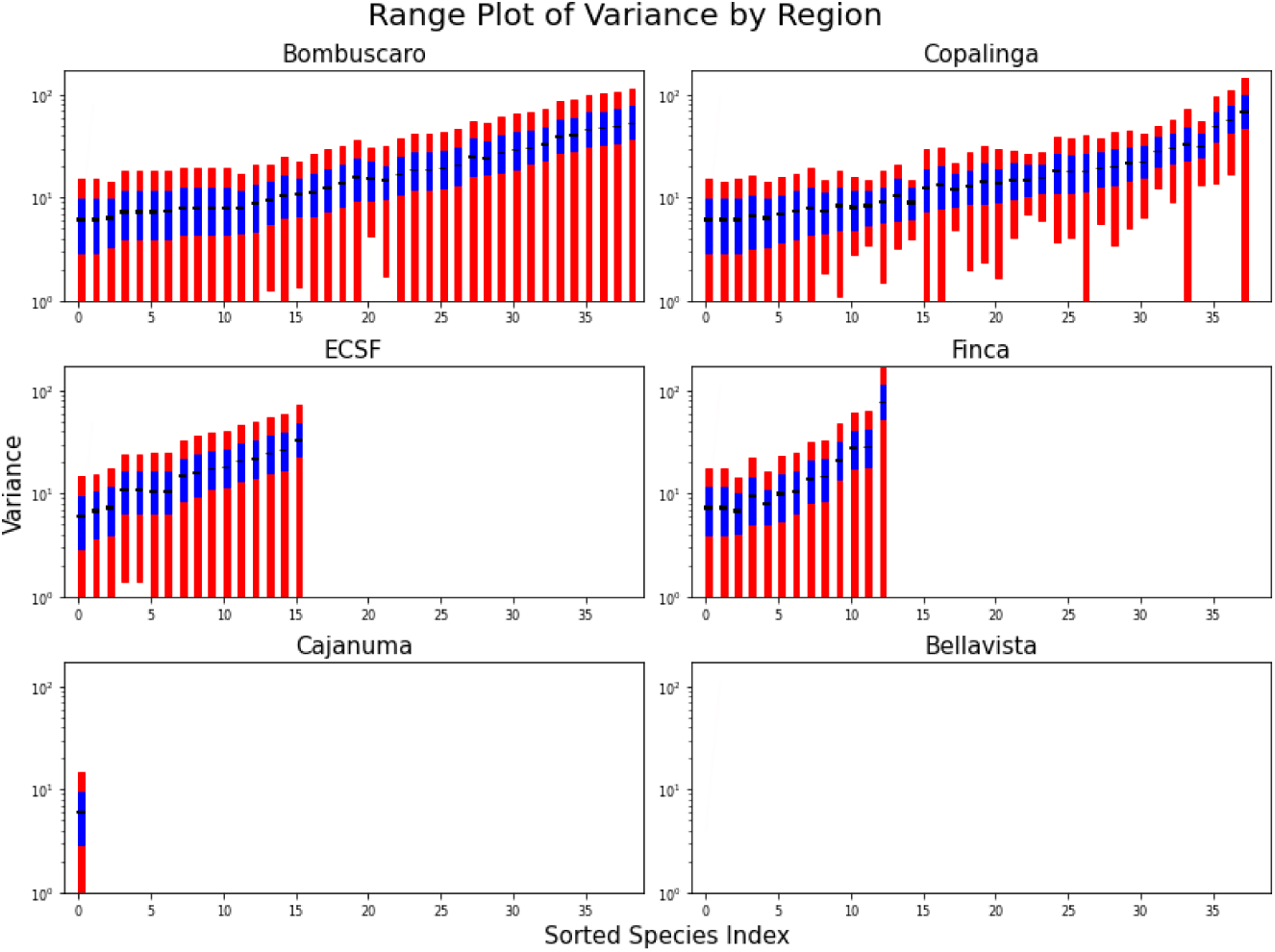
Estimated mean interaction strengths (black segment), 90% confidence intervals for mean interaction strength (blue bars) and 90% confidence interval for location-specific interaction strength (red bar) for different frugivore bird species-fruit species interactions in the six different regions.In each graph, species-species interactions are sorted by increasing mean interaction strength in the given region. (Orderings are not the same for different regions.)

**Figure 8:**
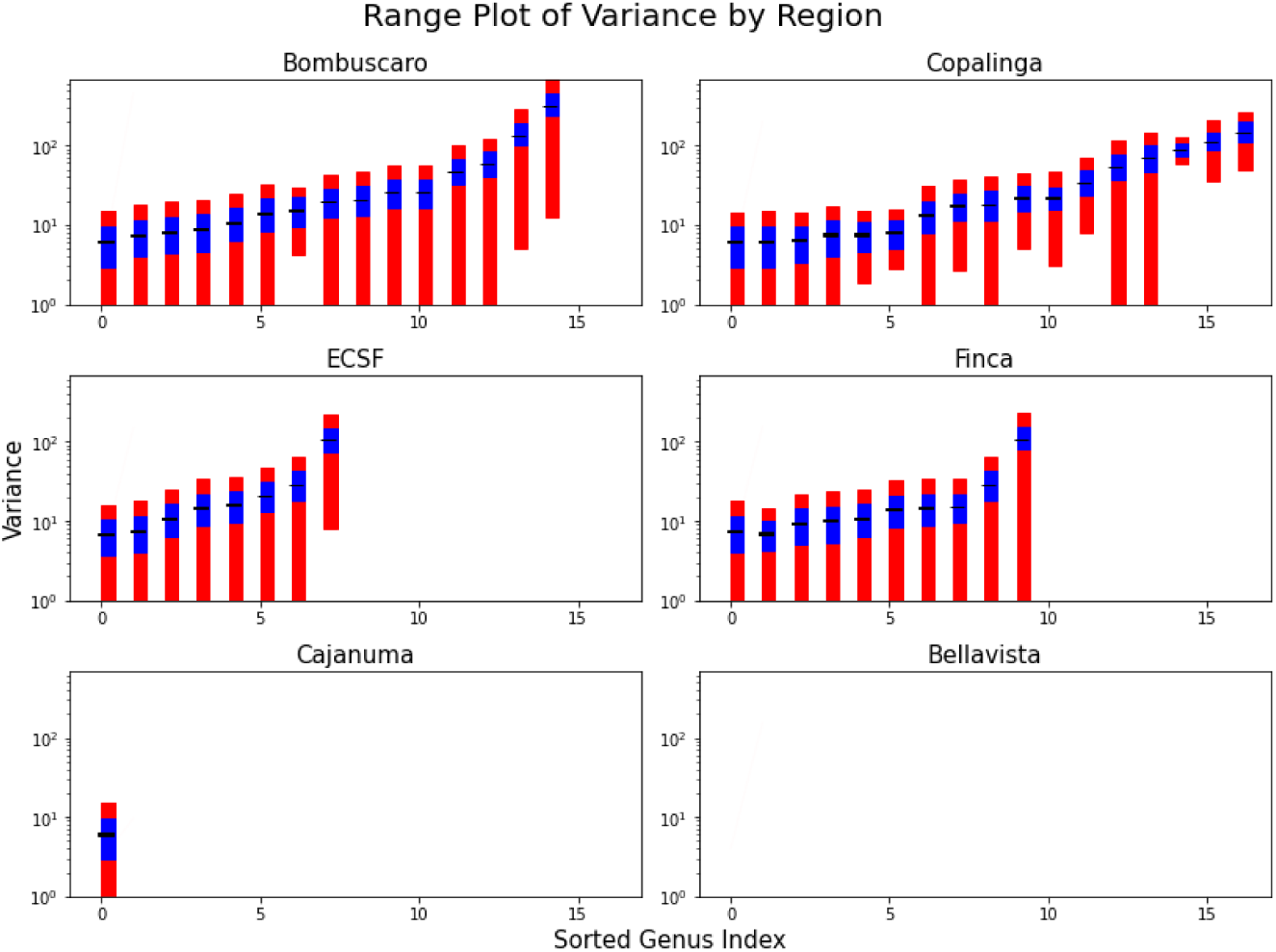
Estimated mean interaction strength (black line), 90% confidence interval for mean interaction strength (blue bar) and 90% confidence interval for location-specific interaction strength (red bar) for different frugivore bird genus-fruit genus interactions in the six different regions. In each graph, genus-genus interactions are sorted by increasing mean interaction strength in the given region. (Orderings may not be the same for different regions.)

**Figure 9:**
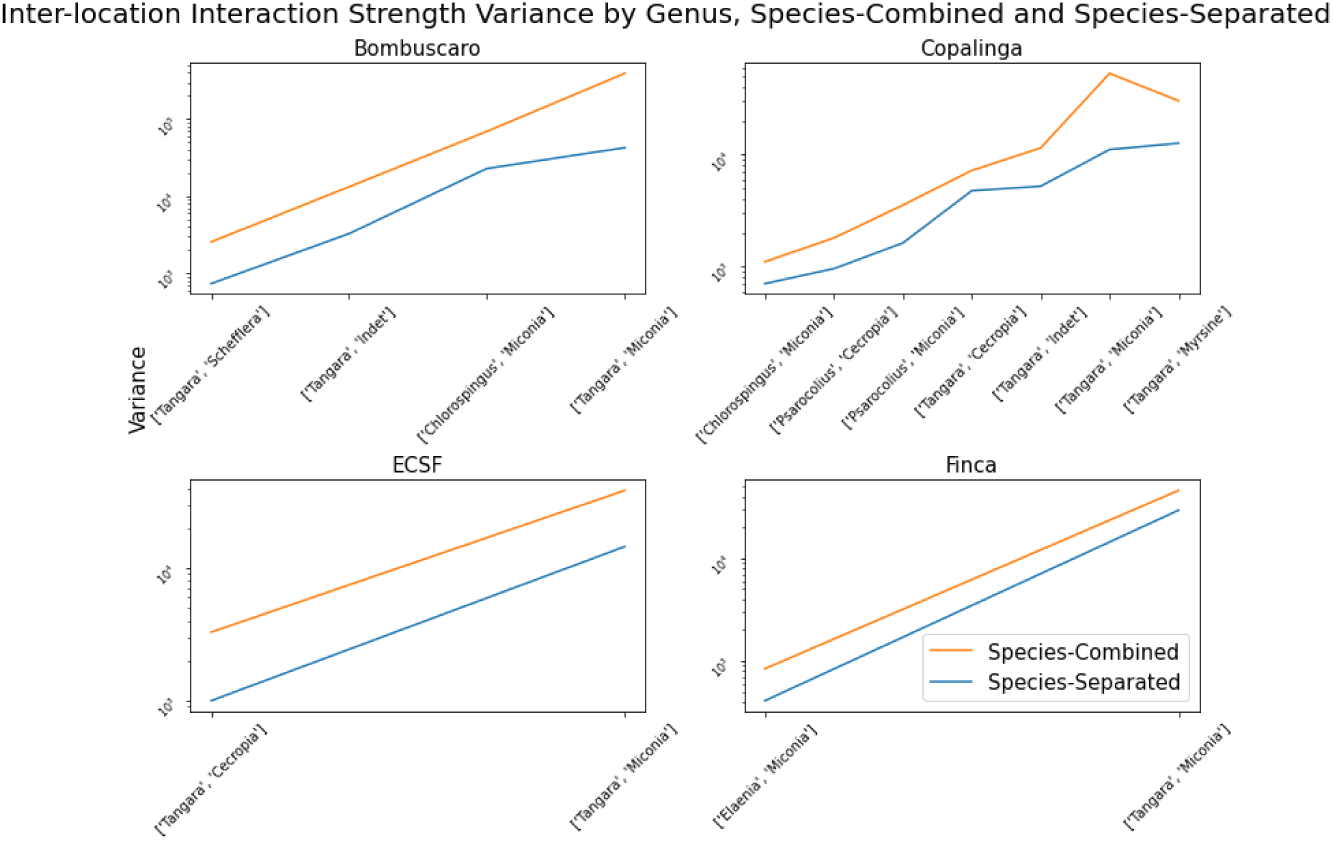
Variance of genus-genus interaction strength (orange) versus sum of variances for all species-species interactions for species included within genus (blue). Species are omitted for which the total number of interactions is less than 10.

**Figure 10:**
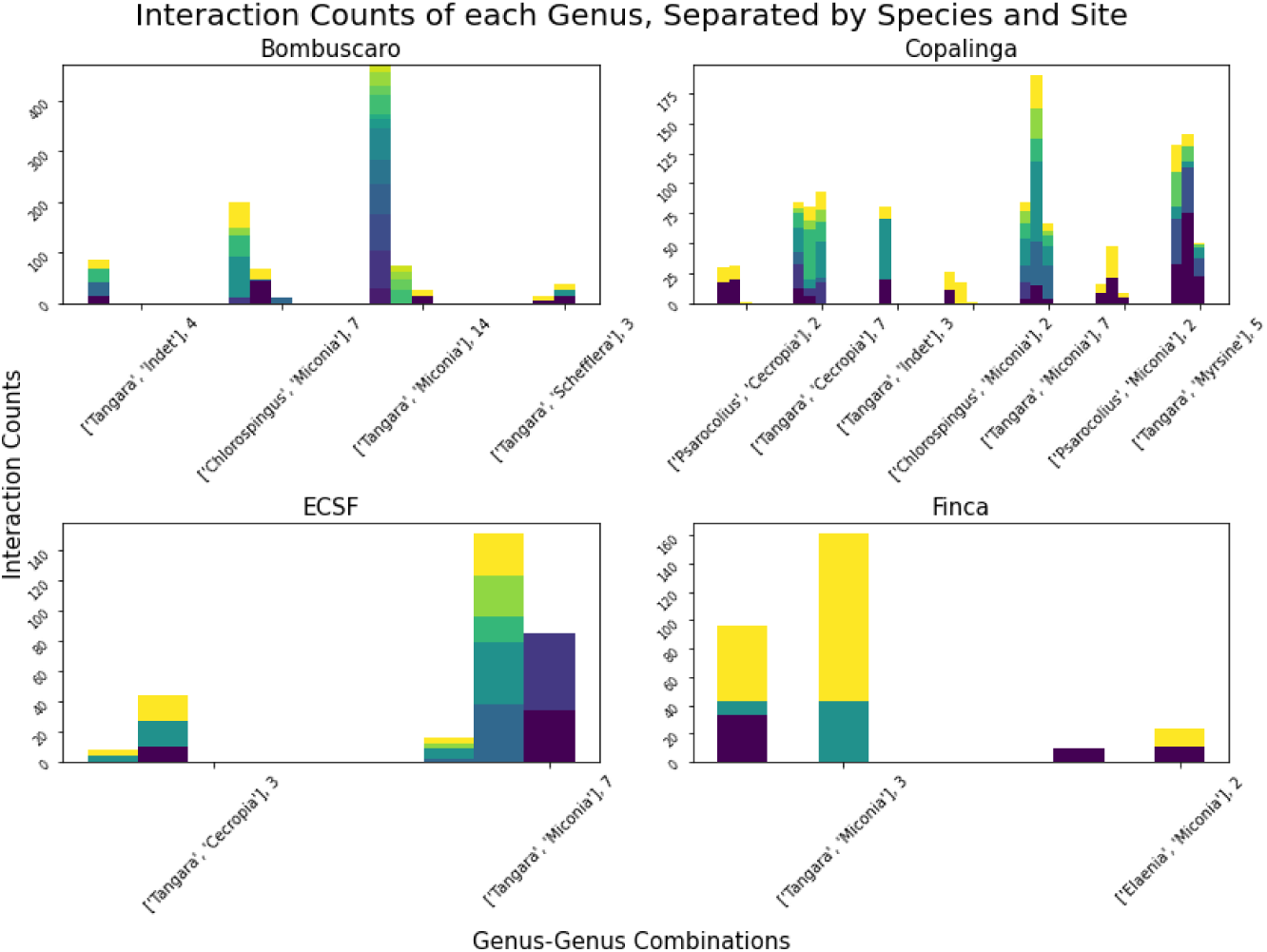
Species-species interaction strengths grouped by genus at 3 locations for each of four regions. (Bellavista and Cajanuma did not have sufficient data.)

## Notes

https://github.com/AaronTAMUCT/Uncertainty

### Competing Interest Statement

The authors have declared no competing interest.

### Summary of Updates

No new data or results were introduced. Background section was added. Paper was reorganized, logic was tightened, Discussion section was expanded, code reference included, AI statement added.

